# Ashwin and FAM98 paralogs define nuclear and cytoplasmic RNA ligase complexes for tRNA biogenesis and the unfolded protein response

**DOI:** 10.1101/2025.08.01.668163

**Authors:** Moritz Leitner, Marius Moser, Nathan Raynaud, Moritz M Pfleiderer, Martin Jinek, Javier Martinez

## Abstract

The tRNA ligase complex (tRNA-LC) seals tRNA exon halves in the nucleus after the removal of a single intron, and joins *XBP1*-mRNA exons in the cytoplasm as part of the unfolded protein response (UPR). This dual function requires simultaneous nuclear and cytoplasmic localization. Here, we reveal that Ashwin (ASW), the vertebrate-specific subunit of the tRNA-LC, serves as its nuclear import factor. ASW displays a dual nuclear localisation signal (NLS) which, upon disruption, leads to the retention of the tRNA-LC in the cytoplasm with a consequent impairment of pre-tRNA splicing and accumulation of 5’ tRNA fragments. We also show that the tRNA-LC exists in three forms depending on which FAM98 paralog is chosen, either FAM98A, FAM98B or FAM98C. We find that ASW interacts exclusively with the FAM98B-containing complex, allowing its nuclear localization for tRNA biogenesis. Attaching an NLS to RTCB, the catalytic and indispensable subunit, rescues pre-tRNA splicing in cells depleted of ASW. We envision that vertebrates evolved ASW to localize a sub-population of tRNA-LC to the nucleus, while using FAM98 paralogs to retain a fraction of RTCB in the cytoplasm for *XBP1*-mRNA splicing during UPR.

## Introduction

Transfer RNAs (tRNAs) are essential adaptor molecules for the translation of messenger RNAs (mRNAs) into proteins. Like most RNAs, tRNAs are synthesized as precursors (pre-tRNAs) that must undergo extensive processing to become functional. One such processing event is splicing, [reviewed in (Gerber *et al*, 2022; Phizicky & Hopper, 2023; Schmidt & Matera, 2019)], a two-step reaction where a single intron is excised by the tRNA splicing endonuclease (TSEN) complex (Paushkin *et al*, 2004; Peebles *et al*, 1983; Trotta *et al*, 1997) and the resulting tRNA exon halves are joined by an RNA ligase activity, generating a mature tRNA (Filipowicz & Shatkin, 1983; Laski *et al*, 1983). While only a subset of eukaryotic pre-tRNA genes contain introns (Chan & Lowe, 2016; Gogakos *et al*, 2017), splicing is essential because for some tRNA families, all iso-decoders contain an intron.

In mammals, the ligation of the tRNA exon halves is catalysed by the tRNA ligase complex (tRNA-LC), composed of the catalytic subunit RTCB, the DEAD-box helicase DDX1, CGI-99 (RTRAF/hCLE), FAM98B, and Ashwin (ASW/C2orf49) (Popow *et al*, 2011; Popow *et al*, 2012). Two additional proteins are essential to sustain the activity of the tRNA-LC: Archease assists the catalytic activity of RTCB by transferring GMP to an essential histidine residue (Gerber *et al*, 2024; Popow *et al*, 2014) and PYROXD1 protects RTCB against oxidative inhibition (Asanovic *et al*, 2021). The tRNA-LC is also required for the splicing of the *XBP1* mRNA during the unfolded protein response, or UPR (Jurkin *et al*, 2014; Kosmaczewski *et al*, 2014; Lu *et al*, 2014). In this event, the *XBP1* mRNA is only incompletely spliced in the nucleus by the spliceosome (*XBP1u* mRNA, u for unspliced), and reaches the cytoplasm with a remaining 26-nt intron that is removed by the endoplasmic reticulum transmembrane endonuclease IRE1 (Sidrauski & Walter, 1997; Yoshida *et al*, 2001), [reviewed in (Acosta-Alvear *et al*, 2024; Walter & Ron, 2011)]. The tRNA-LC joins the resulting mRNA exons to generate the *XBP1s* mRNA (s for spliced) which encodes a transcription factor that activates UPR genes.

The human tRNA-LC can be functionally split into a tetrameric core composed of RTCB, DDX1, FAM98B and CGI-99, and a fifth subunit of unknown function, Ashwin (ASW), shown to assemble into the tRNA-LC by forming a subcomplex with FAM98B and CGI-99 (Kroupova *et al*, 2021). ASW is not required for the activity of the tRNA-LC *in vitro* (Popow *et al*., 2011), but it is important for embryonal development in *X. laevis*, especially in neuronal tissues, by a still unknown mechanism (Patil *et al*, 2006). FAM98B is another subunit whose function remained obscure until recently, when it was reported to disrupt tRNA metabolism via aggregation, triggered by its glycine rich C-terminal sequence (Yang *et al*, 2025). FAM98B has two paralogs in human cells: FAM98A and FAM98C. FAM98B, together with FAM98A, RTCB and DDX1, was found to be involved in the life-cycle of SARS-CoV-2 infections (Kamel *et al*, 2021), and in dendritic RNA transporting granules together with RTCB, DDX1 and CGI-99 (Kanai *et al*, 2004). FAM98C, instead, has not been found as a component of the tRNA-LC but was linked to the dynamics of microtubules (Schou *et al*, 2014) and as a candidate gene for ciliopathies when displaying loss of function mutations (Shaheen *et al*, 2016). It has been reported that protein arginine N-methyltransferase 1 (PRMT1) dimethylates arginine residues in FAM98A (Akter *et al*, 2016) and that both deletion of FAM98A in HeLa cells or expression of a mutant version unable to become dimethylated suppressed cell migration (Wang *et al*, 2023). FAM98A, but not FAM98B, was also reported to connect lysosomes to microtubules by binding to PLHKEM1, which is mutated in osteopetrosis (Fujiwara *et al*, 2016). Finally, FAM98A has been shown to localize in stress granules (Ozeki *et al*, 2019). FAM98 proteins have further been linked to cancer (Akter *et al*, 2017; He *et al*, 2022; Li *et al*, 2019; Liu *et al*, 2021; Rapier-Sharman *et al*, 2022; Zheng *et al*, 2018).

In this report, we identify the function of ASW as the subunit that confers nuclear localization to the tRNA-LC to execute pre-tRNA splicing. The nuclear localization relies on two nuclear localization signals (NLSs) that we dissect at the single amino acid level. ASW specifically binds to FAM98B and therefore only FAM98B-containing tRNA-LC molecules reach the nucleus. Taken together, we have revealed the molecular mechanism evolved to entail the tRNA-LC with a dual subcellular localization, allowing both nuclear and cytoplasmic functions in non-canonical RNA splicing.

## Results

### Paralogs of FAM98B define subpopulations of the human tRNA ligase complex

Curious about the phylogenetic distribution of tRNA-LC subunits, we analysed the occurrence of RTCB, DDX1, CGI-99, FAM98B and ASW in 812 metazoans from an existing orthologous cluster dataset (Kuznetsov *et al*, 2023). While RTCB, DDX1, CGI-99 and FAM98B – core subunits of the tRNA-LC – occur widely throughout metazoans, ASW appears restricted to vertebrates (**Fig. 1A**). We also noticed that, of all considered species, the average copy number of genes in the FAM98 cluster is higher than for any other subunit; this increase is specific to vertebrates, with a median of three FAM98 genes per species, while invertebrates have a median of one (**Fig. 1B**). Our analysis indicates that the subunit ASW emerged in vertebrates and the FAM98 gene evolved concurrently into three paralogs, FAM98A, FAM98B and FAM98C (**Fig. 1C**).

**Figure 1:**
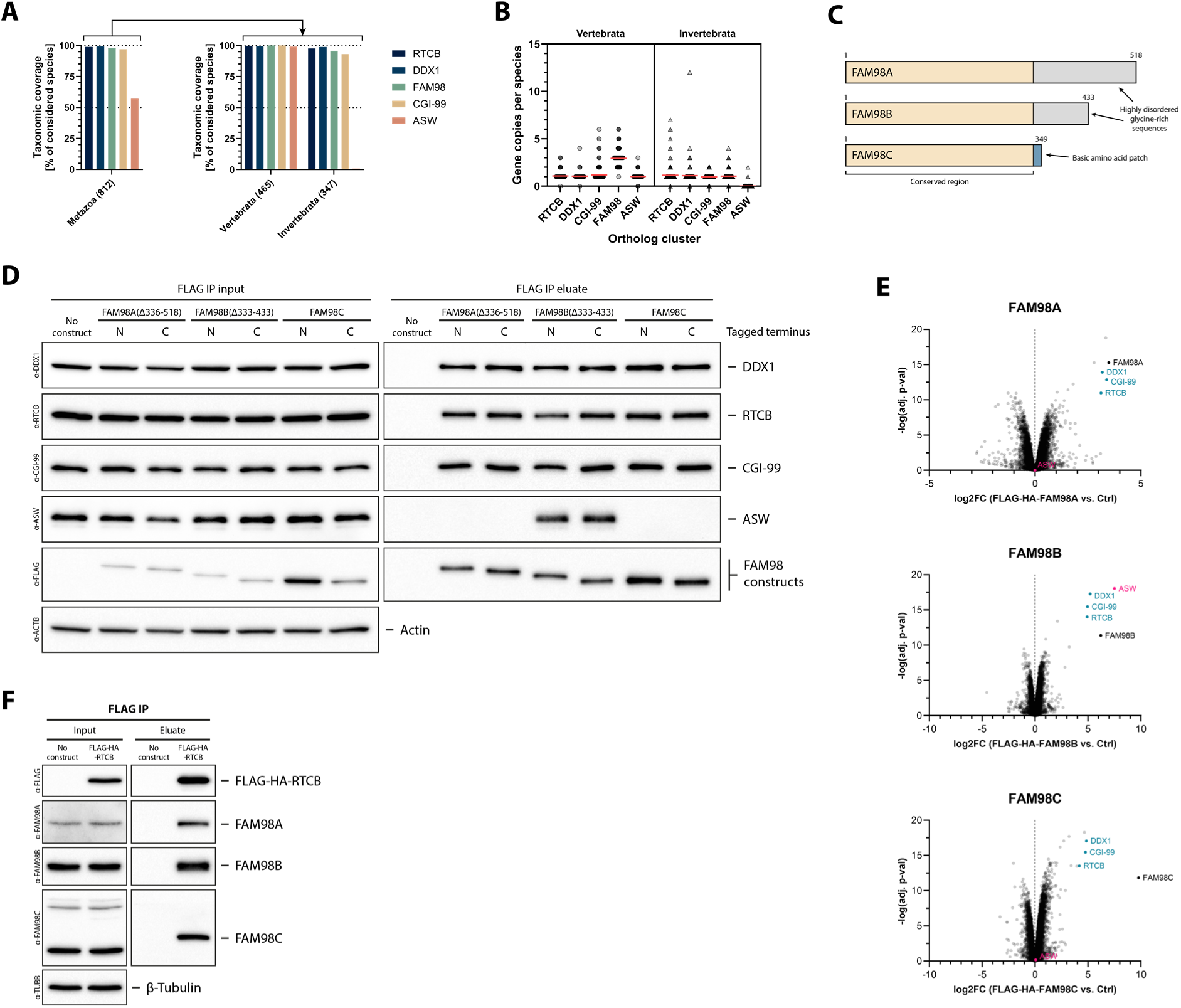
FAM98 paralogs evolved together with ASW in vertebrates to form different populations of tRNA ligase complex. (**A**) Relative number of species having at least one ortholog of RTCB, DDX1, FAM98, CGI-99 or ASW. 812 metazoans were investigated, of which 465 are vertebrates and 347 are invertebrates. Orthologous clusters as well as assigned species were retrieved from OrthoDB v11 (Kuznetsov *et al*., 2023). (**B**) Genes per species that are assigned to the orthologous clusters shown in A. (**C**) Illustration of the three human FAM98 paralogs, highlighting sequences that are conserved or different among them. (**D**) HEK FITR cell lines stably carrying doxycycline-inducible expression constructs for either N– or C-terminally FLAG-HA-tagged FAM98A(Δ336-518), FAM98B(Δ333-433) or FAM98C were treated with doxycycline for 24 h. As control, HEK FITR cells without any integrated construct (no construct) were used. The expressed constructs were isolated using an anti-FLAG IP. Input and eluate samples of the IP were analyzed by western blotting for DDX1, RTCB, CGI-99, ASW, FLAG and ACTB (evaluation of equal loading). Equal amounts of total protein (input) or volume (eluate) were loaded per lane. (**E**) Anti-FLAG IP as in C, but with HEK FITR cells expressing N-terminally FLAG-HA-tagged, full-length FAM98A, FAM98B or FAM98C constructs. After binding and washing, the beads were subjected to tandem mass-spectrometry to identify and quantify bound proteins. The displayed protein enrichments are the results from three independent experiments. (**F**) Anti-FLAG IP as in C, but with HEK FITR cells expressing FLAG-HA-RTCB. Input and eluate samples were analyzed by western blotting for FLAG, FAM98A, FAM98B, FAM98C and ACTB (evaluation of equal loading). Equal amounts of total protein (input) or volume (eluate) were loaded per lane.

To test if all paralogs were able to assemble into the tRNA-LC, we expressed and immunoprecipitated (IP) FLAG-tagged versions of FAM98A, FAM98B and FAM98C from HEK Flp-In T-REx (FITR) cells and performed western blots (WBs) against the remaining subunits of the human tRNA-LC (**Figs. 1D and S1A**). Since FAM98A and FAM98B contain highly repetitive and disordered sequences at the C-terminus, that turned cloning and expression very difficult (Yang *et al*., 2025), we first opted to generate and express truncated versions of FAM98A and FAM98B (**Fig. 1D**). We found all FAM98 paralogs able to associate with RTCB, DDX1 and CGI-99; however, ASW exclusively interacted with both truncated (**Fig. 1D, right panel**) and full-length (**Fig. S1A, right panel**) forms of FAM98B. Mass spectrometry (MS) analysis confirmed that ASW is only enriched in FAM98B IPs (**Figs. 1E and S1B**). We reciprocally detected all three FAM98 paralogs co-eluting with FLAG-tagged RTCB (**Fig. 1F**). Failing to detect a band of the right size for FAM98C (37 kDa) in whole cell lysates (**Fig. 1F, FLAG IP input**) is likely explained by its low abundance, estimated to be around 48 nM in HEK cells (Cho *et al*, 2022) as well as the presence of cross-reacting bands in whole cell lysates. We conclude, that at least three different subpopulations of the tRNA-LC exist in human cells, containing either FAM98A, FAM98B or FAM98C, with ASW exclusively interacting with FAM98B.

### FAM98B and ASW are essential for efficient pre-tRNA splicing

We used CRISPR/Cas9 to delete the genes encoding FAM98A, FAM98B and FAM98C to reveal their role in the stability and enzymatic activity of the tRNA-LC. Deletion of FAM98B destabilized DDX1, RTCB, CGI-99, and more severely ASW (**Fig. 2A**), while no changes were detected upon depletion of FAM98A and FAM98C. Unable to validate the knock-out (KO) of FAM98C through WBs, we turned to MS and IPs of FLAG-HA-RTCB to show efficient deletion of FAM98C (**Fig. S2**). We next tested whether depletion of any FAM98 paralog would impair the ligation of tRNA exon halves during pre-tRNA splicing. We scored a potential defect in ligation via northern blotting (NB), monitoring the accumulation of tRNA fragments composed of 5’ leader and 5’ exon sequences (5’ tRNA fragment), a phenomenon already observed when exposing cells to strong oxidants such as H_2_O_2_ and menadione (Hanada *et al*, 2013). Here, we detected a large accumulation of 5’ tRNA fragments in both FAM98B KO clones, as recently reported by the Chivukula laboratory in HEK293T cells (Yang *et al*., 2025), with no changes in mature, spliced tRNAs (**Fig. 2B, top panels**). 5’ tRNA fragments did not accumulate in FAM98A or FAM98C KO clones (**Fig. 2B, top panels**). As expected, the absence of FAM98B had no impact on non-intron containing tRNAs, tRNA^Met-CAT^ and tRNA^Gly-CCC/GCC^ (**Fig. 2B, bottom panels**). We also found FAM98B dispensable for the splicing of *XBP1* mRNA in the cytoplasm (Yang *et al*., 2025). *XBP1*-mRNA splicing was slightly affected in FAM98A KO cells (**Fig. 2C**) but such defect could not be rescued by re-expression of FAM98A in the FAM98A KO cells, all indicative of a clonal defect in FAM98A KO cells (**Fig. 2D**).

**Figure 2:**
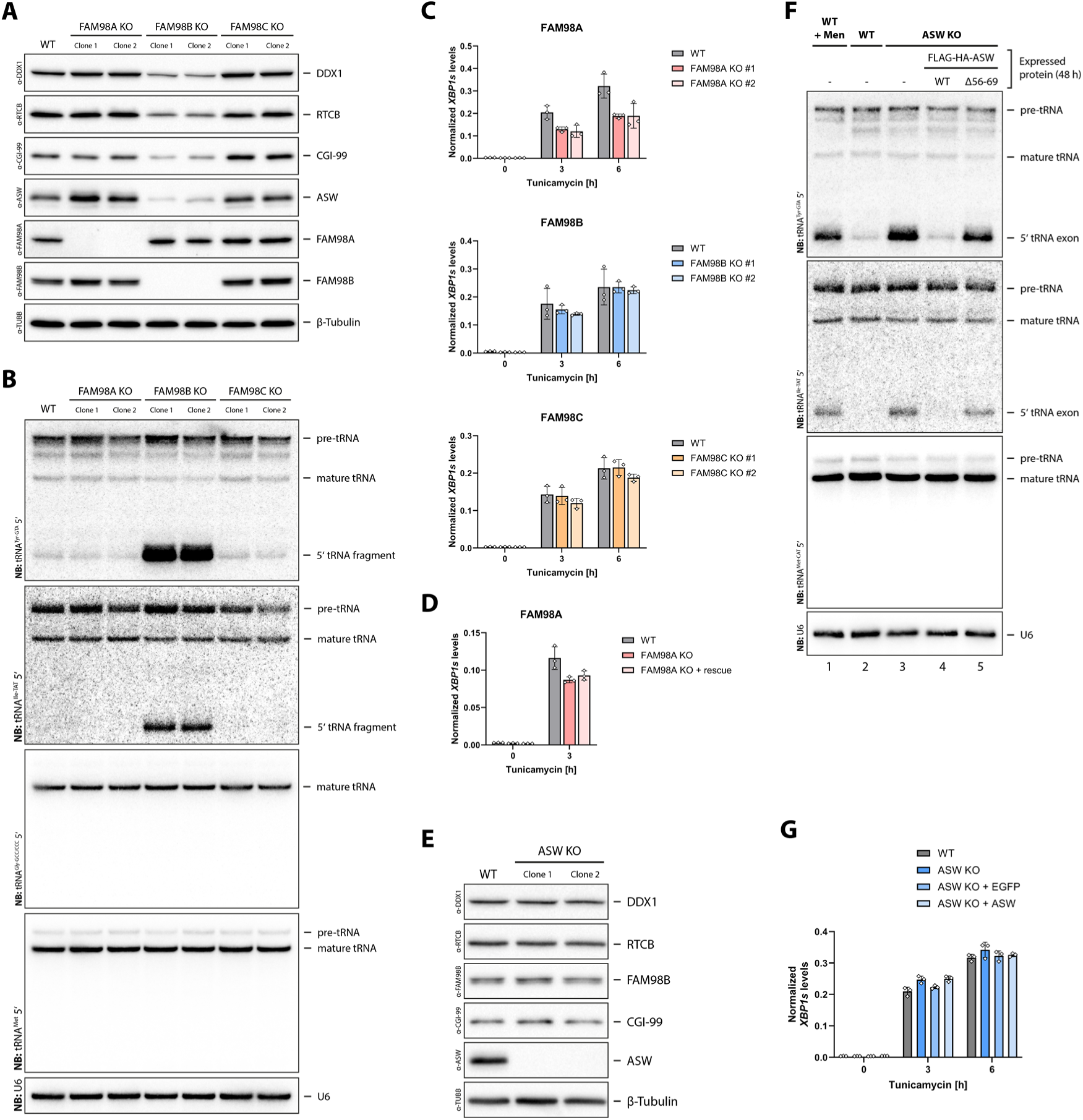
FAM98B and ASW are required for pre-tRNA splicing. (**A**) Whole cell lysates of HEK FITR WT, FAM98A KO, FAM98B KO and FAM98C KO cells were analyzed by western blotting for levels of tRNA-LC subunits as well as TUBB (evaluation of equal loading). Two clones were analyzed for each KO cell line. (**B**) Steady-state levels of tRNA^Tyr-GTA/ATA^, tRNA^Ile-TAT^, tRNA^Gly-GCC/CCC^ and tRNA^Met^ as well as their precursors in HEK FITR WT, FAM98A KO, FAM98B KO and FAM98C KO cells were determined by northern blotting. 5 µg total RNA were loaded per lane. A probe against U6 snRNA was used as loading control. Two clones were analyzed for each KO cell line. (**C**) Relative transcript levels of *XBP1s* (spliced) determined by qPCR during Tunicamycin-induced UPR in HEK FITR WT, FAM98A KO, FAM98B KO and FAM98C KO cells. Expression levels were normalized to *ACTB* mRNA levels. Relative expression levels were calculated with the 2^-ΔΔCt^ method (Livak & Schmittgen, 2001a). Values represent means ± SD from three independent experiments. Two clones were analyzed for each KO cell line. (**D**) UPR experiment as in C, but with HEK FITR WT, HEK FITR FAM98A KO (clone 1) and HEK FITR FAM98A KO (clone 1) cells rescued with FAM98A WT. The rescued FAM98A KO cells have an expression construct for full-length FAM98A stably integrated and expressed. (**E**) Whole cell lysates of HEK FITR WT and ASW KO cells (2 clones) were analyzed by western blotting for levels of tRNA-LC subunits as well as TUBB (evaluation of equal loading). (**F**) Steady-state levels of tRNA^Tyr-GTA/ATA^, tRNA^Ile-TAT^ and tRNA^Met^ as well as their precursors in HEK FITR WT, ASW KO and rescued ASW KO cells were determined by northern blotting. As a positive control for inhibition of the tRNA-LC, WT cells were treated with 40 µM Menadione for 1 h prior to RNA isolation (lane 1). The rescued cells had the recue constructs stably integrated and expressed by the addition of doxycycline. 5 µg total RNA were loaded per lane. A probe against U6 snRNA was used as loading control. (**G**) Same as in C, but with HEK FITR WT, ASW KO and rescued ASW KO cells. The rescued cells had the recue constructs stably integrated and expressed by the addition of doxycycline (1 μg/mL) for 24 h prior to RNA isolation.

With ASW assembling into the tRNA-LC in exclusive association with FAM98B, we wondered whether defective pre-tRNA splicing in FAM98B KO cells was actually due to the concurrent absence of ASW, whose deletion had no impact on the other subunits of the tRNA-LC (**Fig. 2E**) (Popow *et al*., 2011; Popow *et al*., 2014). As in FAM98B KO cells, we also detected accumulation of 5’ tRNA fragments in ASW KO cells, similar to those detected upon inhibition of the tRNA-LC with menadione (**Fig. 2F, lanes 1-3**) (Hanada *et al*., 2013). Overexpression of FLAG-tagged WT ASW rescued pre-tRNA splicing, confirming that the defect is caused by the absence of ASW (**Fig. 2F, lane 4**). However, a mutant of ASW unable to associate with the tRNA-LC due to the deletion of residues 56 to 69 (Δ56-69) (**Fig. S2C**) was not able to abrogate the accumulation of the 5’ tRNA fragment (**Fig. 3B, lane 5**), indicating that ASW supports pre-tRNA splicing via its association with the tRNA-LC. As in FAM98B KO cells, *XBP1*-mRNA splicing remained unaffected in ASW KO cells (**Fig. 2G**).

**Figure 3:**
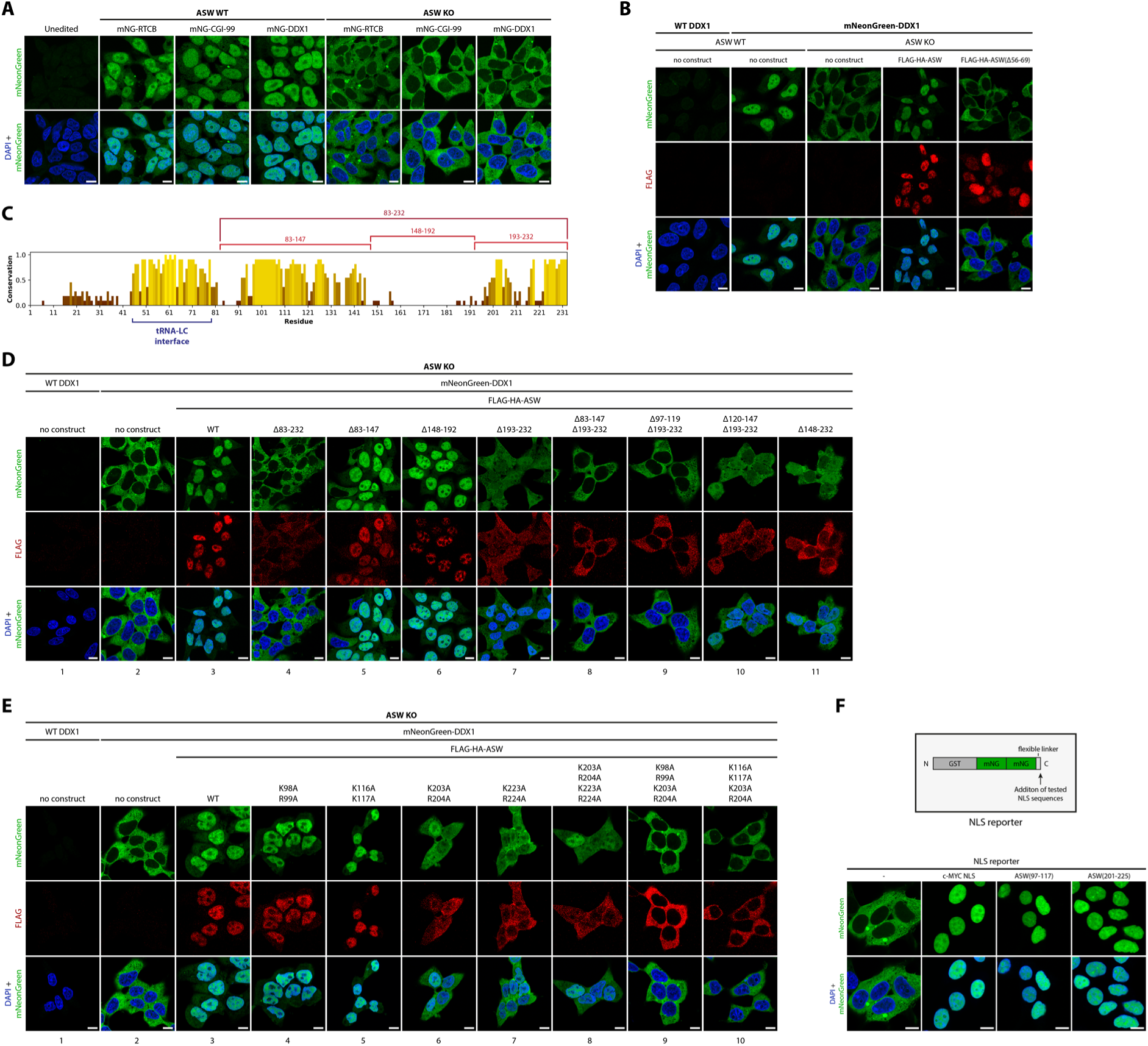
ASW localizes the tRNA-LC to the nucleus using two NLSs that require basic amino acids. (**A**) HEK FITR WT and ASW KO cells expressing endogenous mNeonGreen-RTCB, mNeonGreen-CGI-99 or mNeonGreen-DDX1 were fixed, permeabilized and stained with DAPI, to determine the localization of endogenous tRNA-LC by microscopy. HEK FITR WT cells were used as negative control. (**B**) HEK FITR ASW KO cells endogenously expressing mNeonGreen-DDX1 and stably carrying doxycycline-inducible expression constructs for different FLAG-HA-tagged ASW constructs (WT and Δ56-69) were treated with doxycycline for 24 h. Anti-FLAG immunofluorescence was performed and cells were stained with DAPI, to determine the localization of endogenous tRNA-LC LC as well as the expressed constructs by microscopy. As control, HEK FITR ASW KO cells and HEK FITR ASW KO cells endogenously expressing mNeonGreen-DDX1, without any integrated construct (no construct) were used. (**C**) The conservation of human ASW derived from a multiple sequence alignment (Sievers & Higgins, 2018) of 465 vertebrate ASW sequences. (**D**) HEK FITR ASW KO cells endogenously expressing mNeonGreen-DDX1 and stably carrying doxycycline-inducible expression constructs for different FLAG-HA-tagged ASW constructs (WT and truncation mutants) were treated with doxycycline for 24 h. Anti-FLAG immunofluorescence was performed and cells were stained with DAPI, to determine the localization of endogenous tRNA-LC LC as well as the expressed constructs by microscopy. As control, HEK FITR ASW KO cells and HEK FITR ASW KO cells endogenously expressing mNeonGreen-DDX1, without any integrated construct (no construct) were used. (**E**) Same as in D, but with different point mutants of ASW instead of truncation mutants. (**F**) HEK FITR cell lines stably carrying doxycycline-inducible expression constructs coding for NLS reporters (GST-mNeonGreen-mNeonGreen) with different C-terminally added sequences [c-MYC-NLS, ASW(97-117) or ASW(201-225)] were treated with doxycycline for 24 h. Cells expressing the NLS reporter without any C-terminally added sequence were used as control. The cells were fixed, permeabilized and stained with DAPI, to determine the localizations of the NLS reporters by microscopy. (**A, B, D-F**) Scale bars indicate 10 μm.

### ASW localizes the tRNA ligase complex to the nucleus through two independent nuclear localization signals

Intrigued by the defective pre-tRNA splicing in cells deprived of ASW and the apparent dispensability of ASW for UPR – similarly to cells deprived of FAM98B – we speculated that ASW might play a role in the subcellular localization of the tRNA-LC. Pre-tRNA splicing has been shown to occur in the nucleus of animal cells (De Robertis *et al*, 1981; Melton *et al*, 1980; Paushkin *et al*., 2004) while the UPR-induced splicing of *XBP1* mRNA takes place in the cytoplasm (Gomez-Puerta *et al*, 2022; Yoshida *et al*., 2001). Considering the subcellular locations of these processes, and in hindsight of the reported potential nuclear localization signals (NLSs) in ASW (Patil *et al*., 2006), we hypothesized that ASW might be the protein localizing the tRNA-LC to the nucleus. Thus, we tagged endogenous RTCB, CGI-99 and DDX1 at the N-terminus with mNeonGreen (mNG) (Shaner *et al*, 2013) using CRISPR/Cas9, in both HEK FITR WT and ASW KO cells (**Fig. S3A**). We detected mNG-RTCB, mNG-CGI-99 and mNG-DDX1 mainly in the nucleus but also in the cytoplasm of WT cells (**Fig. 3A**), matching a recent report of mNG-RTCB (Cho *et al*., 2022). In sharp contrast, mNG-RTCB, mNG-CGI-99 and mNG-DDX1 were barely detectable in the nucleus of ASW KO cells and correspondingly accumulated in the cytoplasm, indicating that ASW is required for the nuclear localization of the tRNA-LC. Expression of FLAG-tagged WT ASW, but not ASW Δ56-69, re-imported the tRNA-LC to the nucleus (**Fig. 3B**).

To identify amino acid sequences in ASW responsible for the nuclear localization of the tRNA-LC, we created truncation mutants of FLAG-tagged ASW by removing most of the highly conserved regions (**Fig. 3C**). Expressing those mutants in HEK FITR ASW KO cells endogenously expressing mNG-DDX1 revealed that the NLSs are confined to the last two thirds of the coding sequence of ASW (residues 83-232), downstream of the identified tRNA-LC interface (**Fig. 3D panel 4**). We further split the Δ83-232 mutant into three smaller truncations (**panels 5-7**) and found that only the very C-terminal truncation Δ193-232 (**panel 7**) impaired the localizing function, although not to the extent of the Δ83-232 mutant. We next combined the Δ193-232 mutant with other truncations to identify the remaining part of the NLS. Only a combination of Δ193-232 with Δ83-147 (**panel 8**), that we could further restrict to Δ193-232 with Δ97-119 (**panel 9**) was able to fully abrogate the ability of ASW to localize the tRNA-LC to the nucleus, indicating that ASW harbours functional NLSs only in the regions between residues 97-119 and 193-232 (**Fig. 3D panels 8-11**). We further confirmed that the ASW Δ83-232 mutant, unable to rescue the nuclear localization, is still able to bind the tRNA-LC (**Fig. S3B**).

To further define the NLSs in ASW we used NLS prediction algorithms on the sequence of the protein. The tool NLSEXplorer (Li *et al*, 2025) predicted four NLS segments, all lying within the previously identified regions important for nuclear import (**Fig. S3C**). Assuming that we were dealing with canonical NLSs, consisting of basic amino acid clusters, we mutated arginine and lysine residues into alanine in the predicted NLSs as well as in other highly conserved parts of the central and C-terminal region of ASW (**Fig. S3D**). While most mutants were still able to localize the tRNA-LC efficiently to the nucleus, two mutants in the C-terminal region – K203A R204A and K223A R224A – partially lost such ability (**Figs. 3E panels 6 and 7; and S3E panels 10 and 12**). Correlating with the truncation mutants, those mutants alone were unable to completely abrogate the nuclear localization, neither when combined into a single construct (**Fig. 3E panel 8**). However, combining the K203A R204A mutant with NLS mutants in the 97-117 region – K98A R99A or K116A K117A – resulted in complete loss of nuclear localization (**Fig. 3E panels 9, 10**).

We further tested whether the identified NLSs of ASW can localize an NLS reporter construct (GST-mNG-mNG) to the nucleus and found that both tested regions, ASW 97-117 and 201-225, can independently act as NLS (**Fig. 3F**). We therefore conclude that ASW harbours two functional NLSs, with NLS1 spanning residues 97-117 and NLS2 spanning residues 201-225, and that the function of both NLSs relies on highly conserved basic residues.

### The tRNA ligase complex needs to be localized to the nucleus to execute pre-tRNA splicing

We finally tested whether impairing the nuclear localization of the tRNA-LC by mutating ASW should cause the previously observed defect in pre-tRNA splicing. We overexpressed FLAG-tagged WT ASW as well as truncation and NLS mutants in ASW KO cells and monitored the accumulation of 5’ tRNA fragments (**Figs. 4A and 4B**). All mutants that were unable to re-establish nuclear localization of the tRNA-LC were also unable to abrogate the accumulation of 5’ tRNA fragments (**Compare Figs. 3C and 3D with Figs. 4A and 4B**). Furthermore, ASW mutants that only partially restored the nuclear localization of the tRNA-LC (Δ193-232 in **Fig. 3C**; K203A R204A and K223A R224A in **Fig. 3D**) still nearly completely suppressed the accumulation of 5’ tRNA fragments (**Figs. 4A and 4B**). We performed further rescue experiments in ASW KO cells by overexpressing FLAG-tagged FAM98A, B or C, as well as FLAG-tagged WT RTCB and FLAG-tagged RTCB to which we added the c-MYC NLS (NLS-RTCB) (Dang & Lee, 1988). While overexpression of WT RTCB did not bring the tRNA-LC back the nucleus, overexpression of NLS-RTCB did, although not to a full extent (**Fig. 4C**). Overexpression of FAM98C showed a mild increase in the amount of nuclear tRNA-LC, which was neither observed for FAM98A nor FAM98B. We validated these findings by monitoring the accumulation of 5’ tRNA fragments and found a positive correlation between nuclear exclusion of the tRNA-LC and accumulation of fragments (**Fig. 4C and D**). Overexpression of NLS-RTCB and, to a minor extent, FAM98C, abrogated the accumulation of 5’ tRNA fragments in ASW KO cells almost to a similar extent as when overexpressing WT ASW (Compare **Figs. 4C and D**). Thus, our data indicates that pre-tRNA splicing requires the tRNA-LC to be present in the nucleus. While nature has entitled Ashwin for that function, we show that the tRNA-LC can still fulfil pre-tRNA splicing when reaching the nucleus by other means, like artificially adding an NLS to RTCB or by increasing the levels of the FAM98C-containing tRNA-LC (**Fig. 4D**).

**Figure 4:**
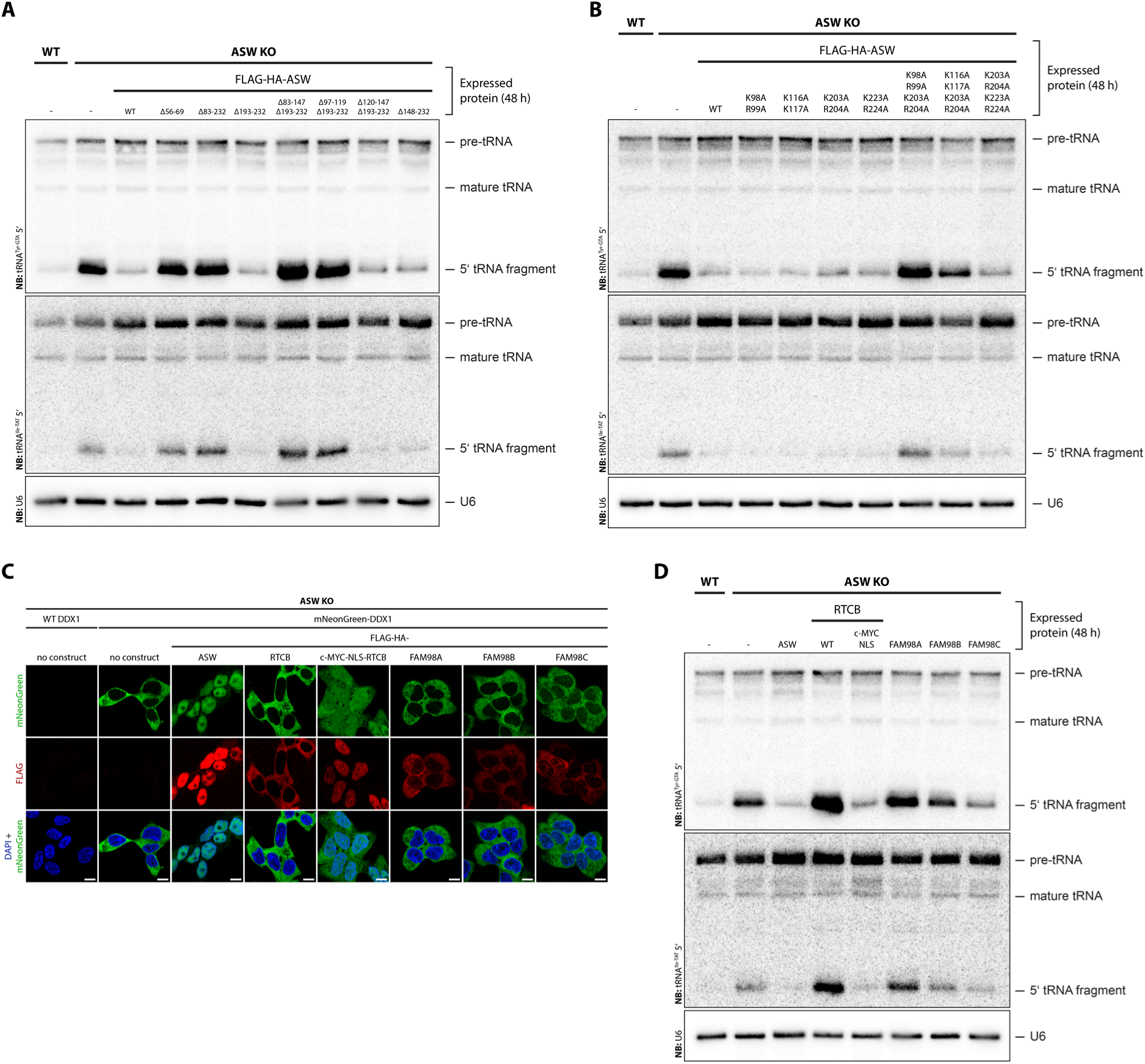
Nuclear localization of the tRNA-LC is essential for efficient pre-tRNA splicing. (**A**) Steady-state levels of tRNA^Tyr-GTA/ATA^ and tRNA^Ile-TAT^ as well as their precursors in HEK FITR WT, ASW KO and rescued ASW KO cells were determined by northern blotting. The rescue cell lines stably carried doxycycline-inducible expression constructs for different FLAG-HA-tagged ASW constructs (WT and truncation mutants) and were treated with doxycycline for 24 h prior to RNA isolation. 5 µg total RNA were loaded per lane. A probe against U6 snRNA was used as loading control. (**B**) Same as in A, but with different point mutants of ASW instead of truncation mutants. (**C**) HEK FITR ASW KO cells endogenously expressing mNeonGreen-DDX1 and stably carrying doxycycline-inducible expression constructs for different FLAG-HA-tagged constructs (ASW, RTCB, c-MYC-NLS-RTCB, FAM98A, FAM98B, FAM98C) were treated with doxycycline for 24 h. Anti-FLAG immunofluorescence was performed and cells were stained with DAPI, to determine the localization of endogenous tRNA-LC as well as the expressed constructs by microscopy. As control, HEK FITR ASW KO cells and HEK FITR ASW KO cells endogenously expressing mNeonGreen-DDX1, without any integrated construct (no construct) were used. Scale bars indicate 10 μm. (**D**) Northern blots as described in A and B, but with the constructs that were used for the microscopy experiments in C.

## Discussion

The advent of vertebrates coincided with the emergence of ASW and the diversification of FAM98 into three paralogs, FAM98A, B and C, leading to sub-populations of tRNA-LC with different subcellular localizations. ASW provides nuclear localization to the tRNA-LC in strict association with FAM98B, thereby allowing pre-tRNA splicing.

The nuclear importing function of ASW relies on two independent NLSs, whose critical residues are highly conserved throughout all analysed ASW homologues. Both NLSs show the characteristics of a bipartite NLS, having two short basic amino acid clusters that are separated by a linker region. While the linker regions are unusually long – 14 and 18 residues – and therefore do not fit the definition of canonical bipartite NLSs (Lu *et al*, 2021), it was shown that bipartite NLSs with longer linker regions exist and can be functional (Lange *et al*, 2010).

We observed severe differences in the expression of human FAM98 paralogs, with FAM98A and FAM98B being less stable than FAM98C. The long and aggregation-prone, glycine-rich tails of FAM98A and, to a larger extent, FAM98B (Yang *et al*., 2025) might dictate this behaviour; removing them equalized their expression. However, considering that FAM98B makes up the majority of the FAM98 proteins in the investigated cell line, the observed effects on stability could have been an artifact caused by overexpression-induced aggregation of FAM98B, which has recently been reported to cause neurodegenerative diseases (Yang *et al*., 2025). Whether the different tails of the FAM98 proteins have any physiological role is yet to be determined.

Removing FAM98B from HEK cells caused a severe co-depletion of all other tRNA-LC subunits, even stronger than previously reported using siRNAs against FAM98B (Popow *et al*., 2011) [see also (Yang *et al*., 2025)]. Structural studies on the assembly of the tRNA-LC showed that FAM98B forms a dimer with CGI-99, which then binds RTCB (Chamera *et al*, 2025; Kroupova *et al*, 2019). We assume that both FAM98A and FAM98C equally assemble into the tRNA-LC, as the CGI-99 and RTCB interfaces are conserved between the FAM98 proteins; in that sense, removal of any FAM98 protein should have the same impact on the stability of the complex. We therefore speculate that the sub-populations of tRNA-LC containing FAM98A and FAM98C are minor in HEK cells and that their depletion would not impact the other subunits in an extent that is detectable by WB. However, this might depend on the cell line, as shRNA-mediated depletion of FAM98A was shown to reduce the levels of other core subunits in A549 cells (Kamel *et al*., 2021).

While we have unveiled a function of the FAM98B/ASW-containing tRNA-LC, the roles of the other tRNA-LC populations remain unclear. Hinted by our taxonomic analysis, we believe that the occurrence of ASW, together with the emergence of several FAM98 paralogs, evolved to support additional functions or regulatory layers for the tRNA-LC, with one being the subcellular distribution of the complex. We believe that splicing of *XBP1* mRNA during the UPR is preserved by keeping certain tRNA-LC subpopulations strictly cytoplasmic, mainly those containing FAM98A. The complementary work of Pfleiderer et al., which dissects the structural framework of human tRNA-LC assembly, supports this idea by showing that FAM98A overexpression leads to a slight defect in pre-tRNA splicing, likely caused by retaining the tRNA-LC in the cytoplasm due to FAM98A out-competing FAM98B for assembly into the tRNA-LC. Beyond the differences in subcellular distribution, FAM98A has for instance been implicated in bone homeostasis (Fujiwara *et al*., 2016) and all three mouse FAM98 paralogs have recently been shown to play roles in osteoclastogenesis (Wang *et al*, 2025). A specialized function of FAM98 paralogs in bone homeostasis would certainly align with their exclusive occurrence in vertebrates. However, which role the tRNA-LC could play in such processes is unknown and would require further investigation.

HEK cells lacking ASW displayed a defect in pre-tRNA splicing, but neither showed a detectable reduction in levels of mature intron-containing tRNAs nor display reduced viability. We believe that a small amount of tRNA-LC might reach the nucleus independently of ASW and provide enough RNA ligase activity to maintain pools of mature tRNAs. In fact, we have seen that a minimal amount of nuclear tRNA-LC is sufficient to rescue the observed splicing defect and we have shown that FAM98C allows the tRNA-LC to reach the nucleus. We therefore suspect that FAM98C might compensate to some extent for the loss of ASW. However, since FAM98C KO cells do not display a defect in pre-tRNA splicing, ASW would still remain as a main driver for the nuclear localization of the tRNA-LC in HEK cells. This might not be true for other cell lines or tissues, where FAM98C might be expressed at higher levels than ASW. Interesting in that context is a certain anticorrelation between ASW and FAM98C expression in some human tissues, as displayed in the human protein atlas dataset (Karlsson *et al*, 2021) hinting that nuclear localization of the tRNA-LC might be achievable in two different ways and that this might be tissue-specific. Contrary to the lack of a strong phenotype in our experiments, it was shown that controlled levels of ASW are essential for cell survival and correct tissue patterning during embryogenesis of *X. laevis* (Patil *et al*., 2006). This suggests that functions of ASW might be specifically needed in the context of tissue formation in multicellular organisms and could thus explain why our simplified tissue culture model would fail to show severe phenotypes in the absence of ASW.

It is indeed fascinating that the nuclear localizing function of the tRNA-LC evolved in the form of a novel subunit and not as a feature in the sequence of FAM98B. We surmise that placing the tRNA-LC at the nucleus is what matters for efficient pre-tRNA splicing in humans, independently of the chosen path. In turn, it is enigmatic how nuclear localization of the tRNA-LC is achieved in invertebrates and whether this is the case at all. While there is previous evidence showing that pre-tRNA splicing in vertebrates takes place in the nucleus (De Robertis *et al*., 1981; Melton *et al*., 1980; Paushkin *et al*., 2004), mechanistic insights into the subcellular localization of this process are lacking for invertebrates, where other mechanisms may allow for nuclear translocation of the tRNA-LC.

Rather surprisingly, we did not observe any decrease in *XBP1*-mRNA splicing in cells lacking either ASW or a FAM98 protein. Loss of DDX1 was recently shown to be innocuous towards *XBP1*-mRNA splicing (Suzuki *et al*, 2025). These observations prompt the idea that *XBP1*-mRNA splicing might require no other subunits than RTCB – or no function provided with the other subunits although being present, excepting Archease and PYROXD1, essential factors for catalysis and protection of the tRNA-LC against oxidative stress (Asanovic *et al*., 2021; Popow *et al*., 2014; Suzuki *et al*., 2025).

RTCB was found to play a role in follicular development; when absent, maternal mRNAs still containing introns accumulated (Zhang et al, 2022). It has also been shown that RTCB, together with AKAP17A and CAAP1, participates in SOS splicing both in human cells and in *C. elegans* (Zhao *et al*, 2025). SOS splicing prevents DNA transposition by acting at the mRNA level, removing transposable levels through excision by a yet unidentified endonuclease, and ligation by RTCB. Based on our results, we hypothesize that SOS splicing should fail in the absence of ASW or show a marked impairment in case FAM98C supports nuclear localization. We also hypothesize that RTCB executes SOS splicing as part of a FAM98B-containing tRNA-LC.

FAM98B was recently shown to aggregate in fragile X-associated tremor/ataxia syndrome (FXTAS) and neuronal intranuclear inclusion disease (NIID), both neurodegenerative disorders linked to expanded GGC-repeats. Those aggregates cause depletion of tRNA-LC and concomitant pre-tRNA splicing defects (Yang *et al*., 2025). While this study highlighted that FAM98B-containing tRNA-LC is nuclear, we provide a mechanistic explanation of how this subpopulation reaches the nucleus and that ASW is ultimately the subunit enabling efficient pre-tRNA splicing due to its nuclear localizing function.

## Materials and Methods

### Cell culture, transfections and UPR-induction

Cells were cultured in complete DMEM [DMEM (Thermo, 41966029) supplemented with 10 % fetal calf serum (Thermo, 10270), 100 U/mL penicillin, 100 μg/mL streptomycin (Lonza, DE17-602E) and 0.2 M HEPES (Thermo, 15630080)] at 37 °C and 5 % CO2. Cells were regularly tested for mycoplasma contamination. For transient transfections, cells were transfected using Lipofectamine 3000 (Thermo, L3000015), following the manufacturer’s instructions. For experiments that required induction of UPR, we incubated the cells with 1.5 µg/mL tunicamycin for 3 or 6 h.

### Taxonomic and conservational analysis

The following orthologous cluster datasets were downloaded from OrthoDB v11 (Kuznetsov *et al*., 2023): 2522645at33208 (RTCB), 125852at33208 (DDX1), 5614at33208 (RTRAF/CGI-99), 7650at33208 (FAM98) and 2482041at33208 (ASW). We also retrieved a list of all metazoans that were analyzed by OrthoDB v11 and further split this list into invertebrates and vertebrates. For calculating the taxonomic coverage of the investigated orthologous clusters, we determined the percentage of species (metazoans, vertebrates or invertebrates) that had at least one gene in the corresponding orthologous cluster. We further determined how many assigned genes each species had per investigated orthologous cluster. For conservational analysis of ASW, we performed a multiple sequence alignment of all protein sequences in the ASW cluster with Clustal Omega (Sievers & Higgins, 2018). From the alignment, we then determined conservation scores for each residue with the software JalView (Waterhouse *et al*, 2009). As we were interested in the conserved residues of human ASW, we omitted all position of the alignment that corresponded to gaps in the sequence of human ASW.

### Construct cloning and mutagenesis

All expression constructs were cloned into the pcDNA5/FRT/TO vector (Thermo, V652020) under a Tet-On-controlled CMV promotor. Constructs were assembled from gel-purified PCR products or from synthesized DNA fragments using NEBuilder HiFi DNA Assembly Master Mix (NEB, E262), according to the manufacturer’s instructions. PCR fragments were amplified using Q5 Hot Start High-Fidelity DNA Polymerase (NEB, M0493), either using cDNA as or existing lab-internal plasmids as template. Primers for site-directed mutagenesis of constructs were designed with NEBaseChanger (https://nebasechanger.neb.com/) and the linear mutagenized constructs were generated with Q5 Hot Start High-Fidelity DNA Polymerase (NEB, M0493) from the non-mutagenized constructs. For circularization of linear mutagenized constructs 2 μL of the unpurified PCR product was mixed with 2 μL 10x rCutSmart (NEB, B6004), 2 μL 10 mM ATP, 1 μL 100 mM DTT, 1 μL T4 DNA Ligase (NEB, M0202), 0.2 μL T4 PNK (NEB, M0201), 0.2 μL DpnI (NEB, R0176) and 11.6 μL nuclease-free water. The circularization reaction was performed in a thermocycler: 15 min at 37 °C, followed by 5 min at 23 °C. Cloning reactions, both from general cloning and mutagenesis, were transformed into competent NEB 5-alpha *E. coli* cells (NEB, C2987) and resulting colonies were further grown as liquid cultures, from which plasmid DNA was isolated. All generated constructs were validated by Sanger sequencing.

### Stable cell line generation (Flp-In T-Rex 293 cells)

To generate stable cell lines, HEK Flp-In T-REx 293 cells (Thermo, R78007) were co-transfected with the pOG44 vector (Thermo, V600520) and a pcDNA5/FRT/TO vector (Thermo, V652020) carrying the construct to be integrated under a Tet-On-controlled CMV promotor. As negative control for the selection, only the pOG44 vector was transfected. Two days after the transfection, the cells were split into selection medium [complete DMEM with 100 µg/mL Hygromycin B (Sigma, H3274)]. The selection was stopped after a minimum of two weeks and when all cells in the negative control were dead. The selected cells were then tested for construct expression upon doxycycline addition (1 μg/mL) and further passaged as usual.

### Cell lysate preparation and immunoprecipitation

Medium was removed, the cells were put on ice and were washed once with ice-cold PBS (Thermo, 14190094). The cells were resuspended by adding ice-cold PBS and pipetting several times up and down. The cells were pelleted at 1000 g, 4 °C for 1 min. The supernatant was completely removed and the pellet was resuspended in a small volume of lysis buffer (30 mM HEPES pH 7.3, 100 mM KCl, 5 mM MgCl_2_, 104 µM AEBSF, 1.0 % NP-40, 10 % Glycerol). The crude lysate was rotated at 18 rpm for 30 min at 4 °C and then pelleted at 21100 g and 4 °C for 10 min. The cleared lysate (supernatant) was transferred into a new tube and the total protein concentration was measured with a Bradford assay (Biorad, 5000006). The lysate was diluted to the desired concentration with lysis buffer. All lysates that belonged to the same experiment were diluted to the same concentration. Anti-FLAG M2 magnetic beads (Sigma, M8823) were equilibrated by washing them three times with 1 mL ice-cold lysis buffer. For binding, the prepared lysates were added to the beads and incubated for 2 h at 4 °C rotating at 18 rpm. For all samples that belonged to the same experiment, equal amounts of beads as well as lysate was used. A small volume of lysate was not added to the beads and kept as input sample. After the binding, the beads were collected with a magnetic separator and the supernatant was removed. The beads were washed five times with 1 mL ice-cold lysis buffer. For elution, ice-cold lysis buffer containing 0.5 mg/mL 3xFLAG peptide was added and mixed by shortly vortexing on the highest setting. The beads were then incubated at 4 °C in a shaker at 1200 rpm for 45 min. The beads were pelleted at 500 g for 1 min at 4 °C and were then collected with a magnetic separator. The eluate was transferred into a new tube. 5x SDS loading dye (5 % SDS w/v, 62.5 mM Tris pH 6.8, 50 % Glycerol v/v, 25 mM EDTA, 0.025 % Bromophenol blue w/v, 5 % β-Mercaptoethanol v/v) was added to the eluates as well as the input samples to a final concentration of 1x. The samples were mixed, boiled at 98 °C for 1 min and then stored at –20 °C until further analysis.

### Western blotting

Samples in 1x SDS loading dye were boiled for 1 min at 98 °C, mixed and then separated by SDS-PAGE. Polyacrylamide gels were blotted onto methanol-activated PVDF membranes (Biorad, 1620175) using Trans-Blot SD Semi-Dry Electrophoretic Transfer Cells (Biorad, 1703940) and Trans-Blot Turbo Transfer Buffer (Biorad, 10026938). Membranes were then blocked in 5 % milk in PBS-T (PBS with 0.05 % Tween-20) for 1 h on a shaker. The blocking solution was removed, diluted primary antibody was added and incubated at 4 °C on a shaker overnight (ON). The antibody solution was removed and the blot was washed three times with 20 mL PBS-T for 10 min at RT. Secondary antibody diluted 1:5000 in 5 % milk PBS-T was added to the membrane and incubated for 1 h at RT on a shaker. The antibody solution was removed and the membrane was washed three times with 20 mL PBS-T for 10 min at RT. The blot was developed with Clarity Western ECL Substrate (Biorad, 1705061) and imaged with a ChemiDoc Touch (Biorad, 1708370). Used antibodies are listed in supplementary tables 2 and 3.

### Co-IP/MS experiments

For the co-IP/MS experiments, the bait constructs were immunoprecipitated as described above, but were washed eight times in detergent-free lysis buffer. The beads, with bound proteins, were resuspended in 30 µL 2 M urea and 50 mM ammonium bicarbonate and were then digested with 150 ng LysC (MS grade, FUJIFILM Wako chemicals) at RT for 90 minutes. The supernatant was transferred to a new tube. The beads were rinsed with 30µL 50mM ammonium bicarbonate and combined with the supernatant. Disulfide bonds were reduced with 2.4 µL of 250 mM DTT for 30 min at RT before adding 2.4 µL of 500 mM iodoacetamide and incubating for 30 min at RT in the dark. The remaining iodoacetamide was quenched with 1.2 µL of 250 mM DTT for 10 min. Proteins were digested with 150 ng trypsin (Trypsin Gold, Promega) in 1.5 µL 50 mM ammonium bicarbonate overnight. The digest was stopped by the addition of trifluoroacetic acid (TFA) to a final concentration of 0.5 %, and the peptides were desalted using C18 Stagetips (Rappsilber *et al*, 2007). LC-MS analysis was performed on an UltiMate 3000 RSLCnano LC system (Thermo Scientific) coupled to a timsTOF HT (Bruker). The system was equipped with a CaptiveSpray ion source (Bruker), and a Butterfly Portfolio Heater (Phoenix S&T). Peptides were loaded onto a trap column (Acclaim PepMap 100 C18 HPLC Column, 5 mm × 0.3 mm, 5 μm particle size, Thermo Scientific) using 0.1 % TFA as mobile phase, and separated on an analytical column (Aurora Ultimate XT C18, 25 cm × 75 µm, 1.7 µm particle size, IonOpticks), applying a linear gradient starting with a mobile phase of 98 % solvent A (0.1 % FA) and 2 % solvent B (80 % acetonitrile, 0.08 % FA), increasing to 35 % solvent B over 60 min at a flow rate of 300 nL/min. The analytical column was heated to 50 °C. The mass spectrometer was operated in data-independent acquisition (DIA) parallel accumulation serial fragmentation (PASEF) mode (Meier *et al*, 2020). MS2 data were acquired with eight PASEF scans per duty cycle, each containing three m/z windows. The m/z window widths were adjusted based on the expected precursor density, covering a total range of 300–1200 m/z. The ion mobility range was set to 0.64-1.42 V*s/cm, and the accumulation and ramp time was set to 100 ms. TIMS elution voltages were calibrated linearly to obtain the reduced ion mobility coefficients (1/K0) using three Agilent ESI-L Tuning Mix ions (m/z 622, 922 and 1222). Collision energy for fragmentation was scaled linearly with precursor mobility (1/K0), ranging from 20 eV (at 1/K0 = 0.6 V*s/cm) to 59 eV at (at 1/K0 = 1.6 V*s/cm). Raw MS data were converted to htrms format using HTRMS converter (Biognosys). Then it was searched with Spectronaut (19.1, Biognosys) in directDIA+ mode against the Uniprot human reference proteome (version 2024_01, www.uniprot.org), as well as a database of most common contaminants (version 2023_03, https://github.com/maxperutzlabs-ms/perutz-ms-contaminants). The search was performed with full trypsin specificity and a maximum of two missed cleavages at a protein and peptide spectrum match false discovery rate of 1 %. Carbamidomethylation of cysteine residues was set as fixed, oxidation of methionine, and N-terminal acetylation as variable modifications. The cross-run normalization was turned off and all other settings were left as default. Computational analysis was performed using Python and the in-house developed Python library MsReport (version 0.0.25, https://zenodo.org/records/16677904). LFQ protein intensities reported by Spectronaut were log2-transformed and normalized across samples using the ModeNormalizer from MsReport. The missing normalized LFQ intensity values were imputed by drawing random values from a normal distribution after filtering out contaminants, proteins with less than 2 peptides and less than 2 quantified values in at least one group. Differences between groups were statistically evaluated using the LIMMA 3.52.1 (Ritchie *et al*, 2015) at 5 % FDR (Benjamini-Hochberg).

### KO cell line generation

For each targeted gene, we selected a pair of gRNA sequences that cut both in the same and one of the first exons. The distances between the cut sites were chosen so that they would cause a frameshift if repaired without the cut-out fragment and without any additional indels. We cloned the two gRNA sequences into two different versions of the pX458 vector (Ran *et al*, 2013), one co-expressing mCherry, the other co-expressing GFP, as described by (Ran *et al*., 2013). Cells were co-transfected with the two sgRNA/Cas9 vectors. Two days after the transfection, the cells were sorted by FACS (BD FACSMelody) for both GFP and mCherry expression. The mCherry– and GFP-positive cells were expanded for 5 days and then single-cell sorted for negative mCherry and GFP expression. The clonal cell lines were expanded for two weeks and then genotyped by PCR. Clones that showed the intended homozygous deletion were subsequently validated by Sanger sequencing to ensure that the wanted frameshift was in place. The KO clones were finally validated by western blotting. As validation of the FAM98C KO was not possible by western blotting, we validated the KO by mass-spec. Used gRNA sequences are listed in supplementary table 1.

### FAM98C KO validation by MS

HEK FITR WT and HEK FITR FAM98C KO cells were grown to 90 % confluency in a 6-well plate. The medium was removed and the cells were carefully washed once with 1 mL ice-cold PBS. The cells were resuspended by adding ice-cold PBS and pipetting several times up and down. The cells were pelleted at 1000 g, 4 °C for 1 min. The supernatant was completely removed and the pellets were lysed in 100 µL 2 % sodium deoxycholate (SDC) and 100 mM Tris-HCl pH 8.8 at 80 °C for 10 min. After cooled down, 1 µL Benzonase was added and kept on ice for 15 min. Then the samples were sonicated using a Bioruptor at high energy for 5 cycles 30/30. After centrifugating at RT 15,000 g for 10 min, the supernatant was transferred to a new tube. 4 times samples volume of cold (–20 °C) 100 % acetone was added to the tube. After thoroughly mixing, the precipitation was done at 4 °C overnight. After centrifugation at 10 °C 15,000 g for 10 min, the supernatant was removed. The pellets were washed with 500 µL cold (–20 °C) 80 % acetone. After centrifugation at 10 °C 15,000 g for 10 min, the supernatant was removed. After completely removing acetone, the pellets were air dried in hood for 5 min. The protein pellets were dissolved in 100 µL 2 % SDC in 100 mM Tris-HCl pH 8.8, at 23 °C shaking 800 rpm for 1 h. Then the samples were sonicated using a Bioruptor at high energy for 3 cycles 30/30. The pellets were disrupted by vortexing and mixing and then incubated at 23 °C shaking 800 rpm for 1 h. After centrifugating at 10 °C 15,000 g for 10 min, the supernatant was transferred to a new tube and the protein concentration was determined with a micro BCA assay (Thermo) following the vendor’s protocol. 50 µg protein in 15 µL were reduced by adding 250 mM DTT to final concentration of 10 mM for 30 min at 50 °C and alkylated by adding 500 mM IAA to final concentration of 20 mM and incubating for 30 min at RT in the dark. The remaining IAA was quenched for 10 min by adding 5 mM DTT. The samples were diluted to 1 % SDC with 15 µL 100 mM Tris-HCl pH 8.8. Proteins were digested by adding 500 ng LysC (MS grade, FUJIFILM Wako chemicals) at RT for 3 h. Then, 1 µg trypsin (Trypsin Gold, Promega; 1 µg/μL solution in 1 mM HCl) was added and incubated at 37 °C overnight. The digest was stopped by the addition of 10 % TFA to reach a final concentration of 2 %. SDC was pelleted by centrifugation at 18,500 g for 10 min. The resulting supernatant was desalted using C18 Stagetips (Rappsilber *et al*., 2007). LC-MS analysis was performed on a Vanquish Neo UHPLC system (Thermo Scientific) coupled to an Orbitrap Exploris 480 mass spectrometer (Thermo Scientific). The system was equipped with a FAIMS Pro interface (Thermo Scientific), a Nanospray Flex ion source (Thermo Scientific), coated emitter tips (PepSep, MSWil), and a Butterfly Portfolio Heater (Phoenix S&T). Peptides were loaded onto a trap column (PepMap Neo C18 5mm × 300 µm, 5 μm particle size, Thermo Scientific) using 0.1 % TFA as mobile phase, and separated on an analytical column (Acclaim PepMap 100 C18 HPLC Column, 50 cm × 75 µm, 2 μm particle size, Thermo Scientific), applying a linear gradient starting with a mobile phase of 98 % solvent A (0.1 % FA) and 2 % solvent B (80 % acetonitrile, 0.08 % FA), increasing to 35 % solvent B over 30 min at a flow rate of 230 nL/min. The analytical column was heated to 30°C. The mass spectrometer was operated in DDA mode. Survey scans were acquired from 350-1500 m/z at CV –42, normalized AGC target of 100 %, resolution of 60000. The PRM parameters – precursor m/z and retention time – were built based on a test run. Precursor of peptides of interest were isolated in a 0.7 m/z window and fragmented with 30 % HCD collision energy. Orbitrap resolution was set to 30000, the normalized AGC target to 100 %. Maximal injection time for target peptides was set to 300 ms. PRM measurements were analysed in Skyline [24.1.1.284, (MacLean *et al*, 2010)]. After loading the list of the 4 PRM targets, the raw data were imported. All peptides and their transitions were validated manually based on retention time, relative ion intensities, and mass accuracy. Three of the most intense non-interfering transitions of the target peptides were selected and their peak areas were exported from Skyline. The summed peak areas were used for peptide quantification.

### Endogenous mNeonGreen-tagging

We selected gRNA sequences that cut in the region surrounding the start codon of the targeted gene and cloned them into the pX330-U6-Chimeric_BB-CBh-hSpCas9-hGem(1/110)-P2A-mCherry vector (modified from (Gutschner *et al*, 2016)) as described by (Ran *et al*., 2013). Repair templates were cloned so that they have the following structure: LeftHomologyArm-mNeonGreen-linker-RightHomologyArm. The homology arms contained the sequences surrounding the gRNA cut site (∼500 bp each), with the 3’ end of the LeftHomologyArm ending just after the start codon of the endogenous locus and the 5’ end of the RightHomologyArm beginning with the first codon after the start codon of the endogenous locus. Cells were co-transfected with the corresponding repair template and the sgRNA/Cas9 vector. Two days after the transfection, the cells were sorted by FACS (BD FACSMelody) for mCherry expression. The mCherry-positive cells were expanded for 5 days and then single-cell sorted for positive mNeonGreen expression and negative mCherry expression. The clonal cell lines were expanded for two weeks and then genotyped by PCR. Clones that had the intended homozygous knock-in were subsequently validated by Sanger sequencing of the insertion locus and by western blotting. Used gRNA sequences are listed in supplementary table 1.

### Northern blotting

Total RNA was isolated from cells using TRIzol (Thermo, 15596018), following the manufacturer’s instructions. The RNA samples were mixed with an equal volume of 2x FA buffer (90 % Formamide, 50 mM EDTA pH 8.0, 0.025 % Bromophenol blue w/v, 0.025 % Xylene cyanol w/v), boiled at 95 °C for 3 min and then separated by 10 % UREA-PAGE (National Diagnostics, EC-833), using 0.5x TBE as running buffer (65 mM Tris pH 7.6, 22.5 mM boric acid, 1.25 mM EDTA). The gels were then blotted onto Amersham Hybond-N+ nylon membranes (Cytiva, RPN303B) at a constant current of 250 mA for 3 h, using 0.5x TBE as transfer buffer. After the transfer, the RNA was UV-crosslinked to the membrane and the membrane was blocked in 30 mL pre-heated hybridization buffer [20 mM sodium phosphate pH 7.2, 75 mM sodium citrate, 750 mM NaCl, 7 % SDS w/v, 3 mg sonicated salmon sperm DNA (Thermo, AM9680)] (Livak & Schmittgen, 2001b) for 1 h, rotating at 50 °C. Radiolabelled DNA probes were added to the hybridization buffer and the membranes were incubated ON rotating at 50 °C. The hybridization solution was discarded and the membranes were washed twice for 1 min in pre-heated (50 °C) low stringency wash buffer (75 mM sodium citrate pH 7.0, 750 mM NaCl, 5 % SDS w/v) and then for 1 min in pre-heated (50 °C) high stringency wash buffer (15 mM sodium citrate pH 7.0, 150 mM NaCl, 1 % SDS w/v). For visualization of the radioactive signal, phosphor screens were exposed to the Northern blots and subsequently imaged using an Amersham Typhoon biomolecular imager. Used DNA probes are listed in supplementary table 4.

### RT-qPCR

cDNA was generated from total RNA using the Maxima First Strand cDNA Synthesis Kit (Thermo, K1672), following the manufacturer’s instructions, and was diluted 1:10 in nuclease-free water before downstream analysis. For quantitative measurement of mRNA levels using RT-qPCR, a master mix was prepared for each set of primers: per reaction 5 µL of 2× GoTaq (Promega, A600A), 0.8 µL primer master mix (5 µM forward primer, 5 µM reverse primer) and 2.2 µL nuclease-free water were mixed. A single reaction, in a well of a 384-well plate, consisted of 2 µL diluted cDNA and 8 µL reaction master mix. The RT-qPCR was performed in a CFX384 Touch thermocycler (Bio-Rad) using the following protocol: 95 °C for 5 min, 95 °C for 10 s, 60 °C for 30 s, plate read, repeat previous 3 steps for 60 cycles, 95 °C for 10 s, melt curve determination from 55 °C to 95 °C in 0.5 °C increments. The quality of the PCR products was evaluated by melting curve analysis. Relative expression levels were calculated from Ct values with the 2^-ΔΔCt^ method, normalizing expression levels to *ACTB* mRNA levels (Livak & Schmittgen, 2001b). Used primer sequences are listed in supplementary table 4.

### Immunofluorescence and confocal microscopy

UV-sterilized, quadratic, 12 mm #1.5 coverslips were transferred into a 12-well plate, coated with poly-L-lysine (Sigma, P6282) and seeded with 50-100 k HEK cells. Optionally, constructs for imaging were either transiently transfected or, if using stable cell lines, induced by addition of doxycycline (1 μg/mL). Transiently transfected cells were further processed 48 h after the transfection. Cells that were treated with doxycycline were further processed 24 h after doxycycline addition. The medium was removed and the cells were carefully washed twice with 500 µL PBS. For fixation, 500 µL 4 % paraformaldehyde in PBS was added per well and incubated 15 min at RT. The solution was removed and the cells were washed once with 500 µL PBS. For permeabilization, 500 µL 0.3 % Triton X-100 in PBS was added per well and incubated 5 min at RT. The solution was removed and the cells were washed once with 500 µL PBS. The coverslips were blocked with 50 µL blocking solution (5 % normal goat serum in PBS) for 45 min at RT in the dark. The blocking solution was removed and 30 µL primary antibody diluted in blocking solution was added and incubated for 1 h at RT in the dark. The antibody solution was removed and the coverslips were washed twice with 100 µL PBS. 60 µL secondary antibody diluted 1:1000 in blocking solution was added and incubated for 45 min at RT in the dark. The solution was removed and the coverslips were washed once with 100 µL PBS. 80 µL of 0.5 µg/mL 4’,6-diamidino-2-phenylindole (DAPI) in PBS was added and incubated 10 min at RT in the dark. The solution was removed and the coverslips were washed twice with 100 µL PBS. Finally, the coverslips were mounted onto glass slides with VECTASHIELD PLUS mounting media (Vector Laboratories, H-1900) and sealed with clear nail polish. The slides were stored at 4 °C in the dark until imaging. Images were acquired using an LSM 980 Airyscan 2 confocal microscope (Zeiss). For all samples that belonged to the same experiment, the same laser power settings as well as photomultiplier voltages were used. Used antibodies are listed in supplementary tables 2 and 3.

## Notes

### Competing Interest Statement

The authors have declared no competing interest.

