## Supplementary data for "Ashwin and FAM98 paralogs define nuclear and cytoplasmic RNA ligase complexes for tRNA biogenesis and the unfolded protein response"

### Supplementary Figures

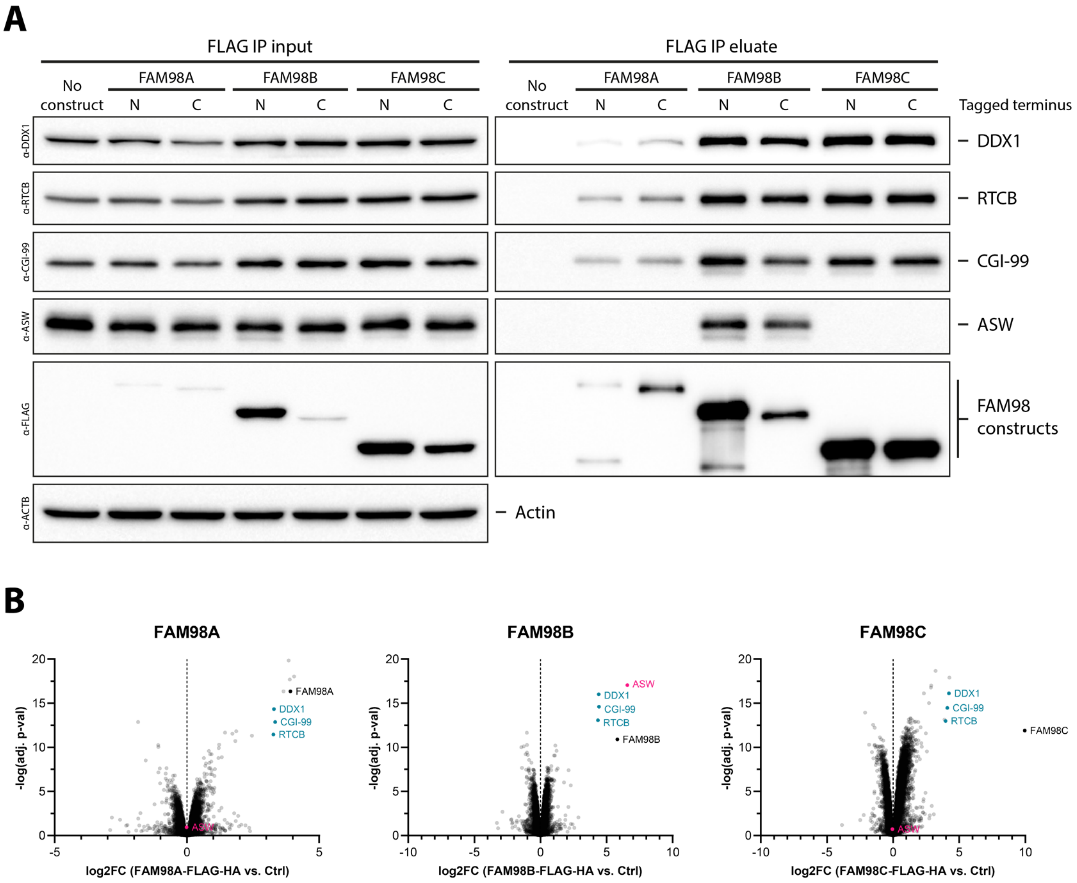


**Figure S1: Only FAM98B-containing tRNA-LC can bind ASW.** (**A**) HEK FITR cell lines stably carrying doxycycline-inducible expression constructs for either N- or C-terminally FLAG-HA-tagged FAM98A, FAM98B or FAM98C were treated with doxycycline for 24 h. As control, HEK FITR cells without any integrated construct (no construct) were used. The expressed constructs were isolated using an anti-FLAG IP. Input and eluate samples of the IP were analyzed by western blotting for DDX1, RTCB, CGI-99, ASW, FLAG and ACTB (evaluation of equal loading). Equal amounts of total protein (input) or volume (eluate) were loaded per lane. (**B**) Anti-FLAG IP as in A, but with HEK FITR cells expressing C-terminally FLAG-HA-tagged FAM98A, FAM98B or FAM98C constructs. After binding and washing, the beads were subjected to tandem mass-spec to identify and quantify bound proteins. The displayed protein enrichments are the results from three independent experiments.

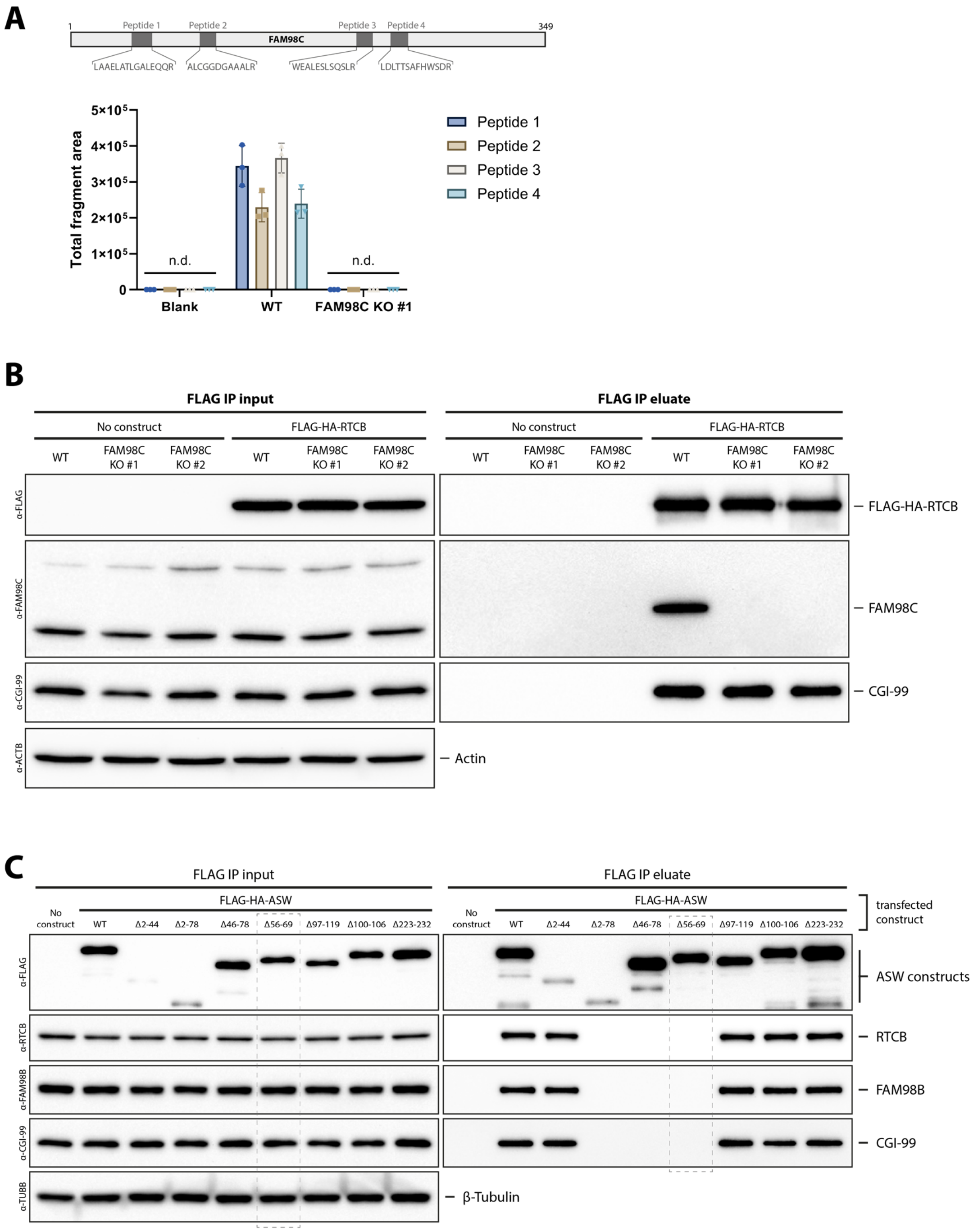


**Figure S2: Validation of the FAM98C KO and ASW tRNA-LC interface mutants.** (**A**) The presence of four FAM98C-specific peptides (see scheme) was determined by tandem mass-spectrometry in cell lysates of HEK FITR WT and HEK FITR FAM98C KO (clone 1) cells. Lysis buffer was used as negative control. Data from three independent experiments. (**B**) HEK FITR WT and HEK FITR FAM98C KO cells were transiently transfected with an expression construct coding for FLAG-HA-RTCB. As control, cells without any transfected construct (no construct) were used. The expressed constructs were isolated using an anti-FLAG IP. Input and eluate samples of the IP were analyzed by western blotting for FLAG, FAM98C, CGI-99 and ACTB (evaluation of equal loading), to validate the KO of FAM98C. Equal amounts of total protein (input) or volume (eluate) were loaded per lane. (**C**) HEK FITR WT cells were transiently transfected with different FLAG-HA-ASW expression constructs (WT and truncation mutants). As control, cells without any transfected construct (no construct) were used. The expressed constructs were isolated using an anti-FLAG IP. Input and eluate samples of the IP were analyzed by western blotting for FLAG, RTCB, FAM98B, CGI-99 and TUBB (evaluation of equal loading), to identify mutants that do not bind the tRNA-LC. Equal amounts of total protein (input) or volume (eluate) were loaded per lane.

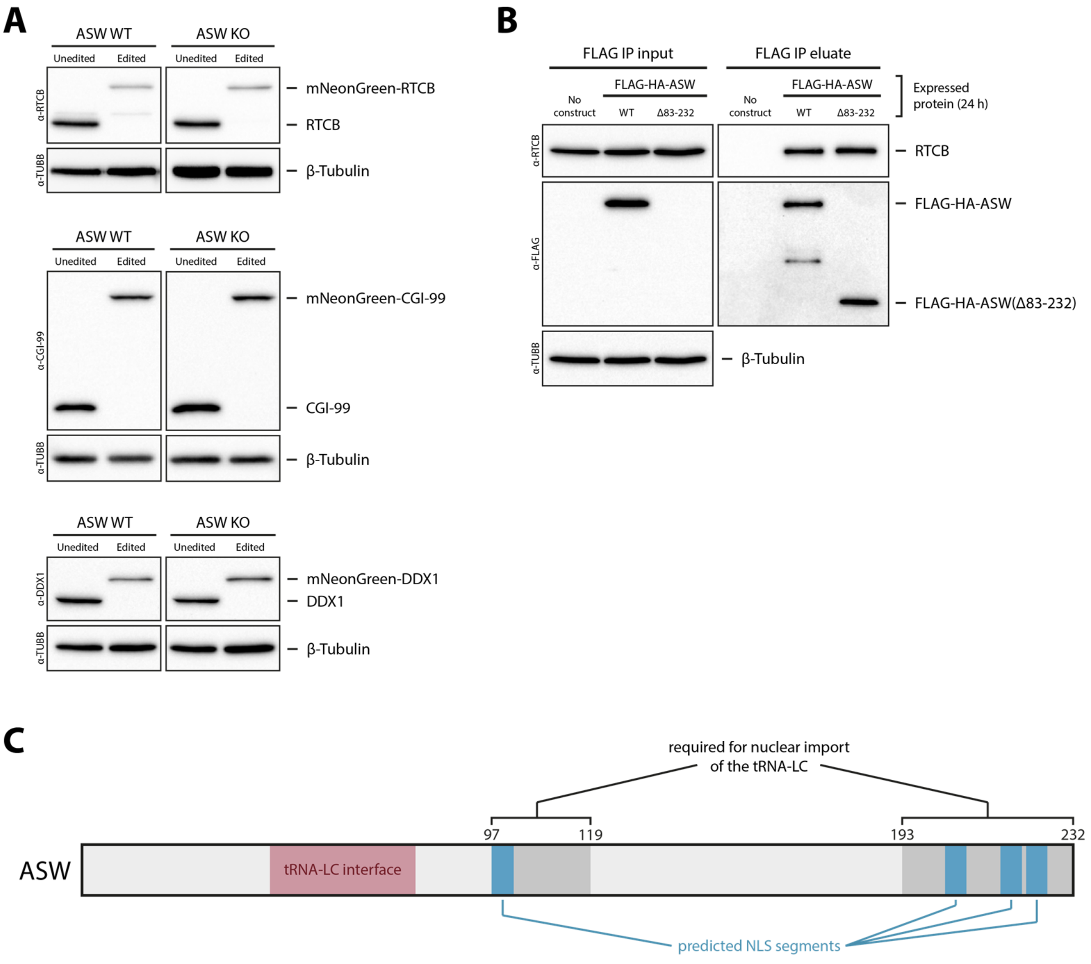


| Position | Recommendation score | Entropy score | Recommended NLS segment |
|---|---|---|---|
| 216-220 | 0.97955 | 1.92193 | NLKPP |
| 97-101 | 0.95852 | 1.92193 | RKRPL |
| 222-226 | 0.94971 | 1.92193 | AKRKI |
| 203-207 | 0.94006 | 1.92193 | KRAAP |

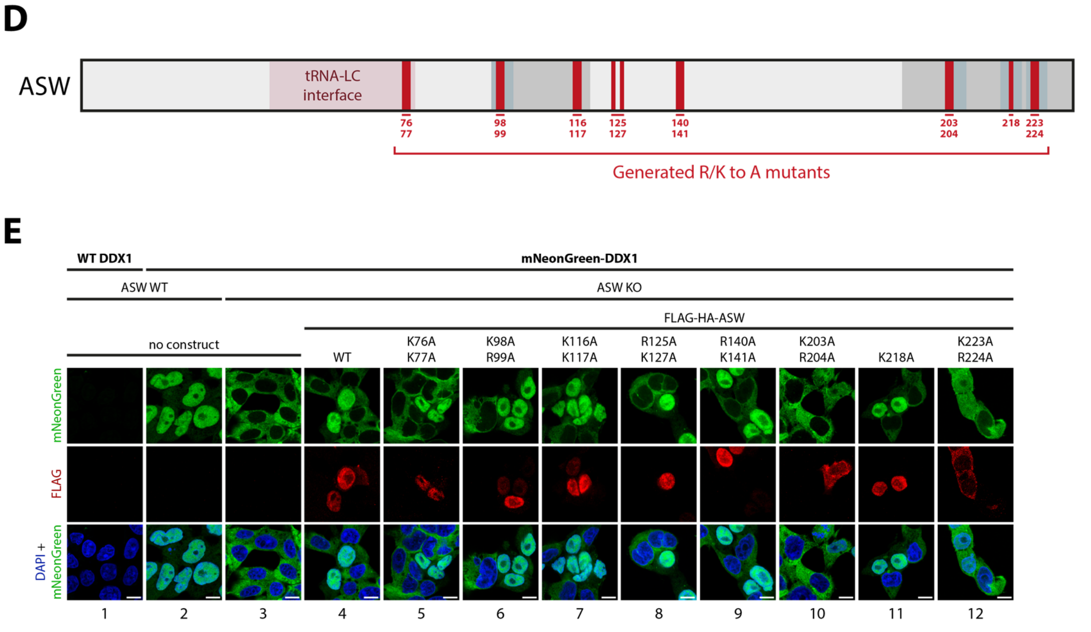

**Figure S3: Validation of the endogenous mNeonGreen knock-ins on RTCB, CGI-99 and DDX1 as well as ASW NLS point mutants.** (**A**) Whole cell lysates of HEK FITR WT and ASW KO cells endogenously expressing mNeonGreen-RTCB, mNeonGreen-CGI-99 or mNeonGreen-DDX1 were evaluated by western blotting for levels of the edited protein as well as TUBB. Equal amounts of total protein were loaded per lane. (**B**) HEK FITR WT cells were transiently transfected with different FLAG-HA-ASW expression constructs (WT and Δ83-232). As control, cells without any transfected construct (no construct) were used. The expressed constructs were isolated using an anti-FLAG IP. Input and eluate samples of the IP were analyzed by western blotting for FLAG, RTCB and TUBB (evaluation of equal loading), to validate that the Δ83-232 ASW mutant still binds the tRNA-LC. Equal amounts of total protein (input) or volume (eluate) were loaded per lane. (**C**) NLS segments in ASW that were predicted by the tool NLSExplorer (Li *et al*, 2025) overlaid on regions of ASW that we identified to contain an NLS. (**D**) Overview of R/K to A mutants that we generated and tested for inactivating the NLSs in ASW. (**E**) HEK FITR ASW KO cells expressing endogenous mNeonGreen-DDX1 were transiently transfected with different FLAG-HA-ASW rescue constructs (WT or R/K to A mutants). Anti-FLAG immunofluorescence was performed and cells were stained with DAPI, to determine the localization of endogenous tRNA-LC LC as well as the expressed constructs by microscopy. As control, HEK FITR WT and ASW KO cells endogenously expressing mNeonGreen-DDX1, without any transfected construct (no construct) were used. Scale bars indicate 10 μm.

### Supplementary Tables

***Supplementary Table 1: Used gRNA sequences.***

| Name | Used for | Sequence |
|---|---|---|
| gRNA_ASW_KO_pair1_a | ASW KO | ATTTGCCGAAGAATAGATGG |
| gRNA_ASW_KO_pair1_b | | GAGGCAATGGTATTGCATGT |
| gRNA_FAM98A_KO_pair1_a | FAM98A KO | AAGGGCCCATTGTTGGAAGA |
| gRNA_FAM98A_KO_pair1_b | | ACAGAGTTTGGTAAACTCGG |
| gRNA_FAM98B_KO_pair1_a | FAM98B KO | AGGGCTTGCTCTTCTAACAA |
| gRNA_FAM98B_KO_pair1_b | | CTTACAAAGGCGGCAGAGGG |
| gRNA_FAM98C_KO_pair1_a | FAM98C KO | ACTTCAGGGGGCTGTGCGTG |
| gRNA_FAM98C_KO_pair1_b | | CGCCTCTCGCTGCTGCTCGA |
| gRNA_RTCB_N-term | mNeonGreen-tagging RTCB | ATTATAGCTGCGACTCATGG |
| gRNA_DDX1_N-term | mNeonGreen-tagging DDX1 | TTCCACAAACGCACCGGAGA |
| gRNA_CGI-99_N-term | mNeonGreen-tagging CGI-99 | CGTCAACTTGCGTCGGAACA |

***Supplementary Table 2: Used primary antibodies.***

| Antibody | Manufacturer | Catalog number | Used for |
|---|---|---|---|
| anti-FLAG | Sigma Aldrich | F1804 | WB, immunofluorescence |
| anti-RTCB | Bethyl Laboratories | A305-079A | WB |
| anti-DDX1 | Bethyl Laboratories | A300-521A | WB |
| anti-FAM98A | Abcam | ab204083 | WB |
| anti-FAM98B | Thermo | PA5-52568 | WB |
| anti-FAM98C | Thermo | PA5-59356 | WB |
| anti-CGI-99 | Abcam | ab188326 | WB |
| anti-ASW | self-made | - | WB |
| anti-TUBB | Abcam | ab44928 | WB |
| anti-ACTB | Sigma Aldrich | A2066 | WB |

***Supplementary Table 3: Used secondary antibodies.***

| Antibody | Manufacturer | Catalog number | Used for |
|---|---|---|---|
| anti-mouse Alexa Fluor Plus 647 | Invitrogen | A32728 | Immunofluorescence |
| anti-rabbit IgG (H+L), HRP | Invitrogen | 65-6120 | WB |
| anti-mouse IgG (H+L), HRP | Invitrogen | 62-6520 | WB |

***Supplementary Table 4: Used Northern probes and RT-qPCR primers.***

| Name | Used for | Sequence |
|---|---|---|
| NB_probe_Tyr_5p | Northern blotting | CTACAGTCCTCCGCTCTACC |
| NB_probe_Ile_5p | Northern blotting | TATAAGTACCGCGCGCTAAC |
| NB_probe_Gly_5p | Northern blotting | TACCACTGAACCACCAATGC |
| NB_probe_Met _5p | Northern blotting | GGGCCCAGCACGCTTCCGCTGCGCCACTCTGC |
| NB_probe_U6 | Northern blotting | GCAGGGGCCACGCTAATCTTCTCTG |
| q_ACTB_FWD | RT-qPCR | GCAGAAGGAGATCACTGCCC |
| q_ACTB_REV | RT-qPCR | GTACTTGCGCTCAGGAGGAG |
| q_XBP1u_FWD | RT-qPCR | ACTACGTGCACCTCTGCAG |
| q_XBP1u_REV | RT-qPCR | GGAAGGGCATTTGAAGAACA |
| q_XBP1s_FWD | RT-qPCR | GAGTCCGCAGCAGGTG |
| q_XBP1s_REV | RT-qPCR | GGAAGGGCATTTGAAGAACA |
| q_XBP1total_FWD | RT-qPCR | GCGCTGAGGAGGAAACTGAAAAAC |
| q_XBP1total_REV | RT-qPCR | CCAAGCGCTGTCTTAACTCC |
